# Chemical-LTP induces confinment of BDNF mRNA under dendritic spines and BDNF protein accumulation inside the spines

**DOI:** 10.1101/2023.10.17.562692

**Authors:** Giorgia Bimbi, Enrico Tongiorgi

## Abstract

The neurotrophin brain-derived neurotrophic factor (BDNF) plays a key role in neuronal development and synaptic plasticity. The discovery that BDNF mRNA can be transported in neuronal dendrites in an activity-dependent manner has suggested that its local translation may support synapse maturation and plasticity. However, a clear demonstration that BDNF mRNA is locally transported and translated at activated synapses in response to long-term potentiation (LTP) is still lacking. Here, we study the dynamics of BDNF mRNA dendritic trafficking following induction of chemical-LTP (cLTP). Dendritic transport of BDNF transcripts was analysed using the MS2 system for mRNA visualization, and chimeric BDNF-GFP constructs were used to monitor protein synthesis in living neurons. We found that within 15’ following cLTP induction, most BDNF mRNA granules become stationary and transiently accumulate in the dendritic shaft at the basis of the spines similarly to the control CamkIIα mRNA which increased also inside the spines, at 60’ post-cLTP. At 60’ but not at 15’ from cLTP induction, we observed an increase in BDNF protein levels within the spine. Taken together, these findings suggest that BDNF mRNA trafficking is arrested in the early phase of cLTP, providing a local source of mRNA for translation of BDNF at the basis of the spine followed in the late LTP phase, by translocation of the BDNF protein within the spine head.

**Statement:** Brain-derived neurotrophic factor (BDNF) plays a key role in neuronal development and synaptic plasticity. In this study, we investigate two unresolved questions in neuronal plasticity: a) whether the post-synaptically released BDNF can be locally synthesized in this compartment, and b) whether the local translation of BDNF occurs in dendrites, or within the spine. Using chimeric constructs ectopically expressed in living primary hippocampal neurons, we tracked BDNF mRNA trafficking within the dendrites and its local translation following induction of chemical-LTP (cLTP) by forskolin. We show that in the early phase of cLTP induction (15’), BDNF mRNA becomes confined at the basis of the spines providing a local source for translation of the protein followed in the late LTP phase (60’), by translocation of the BDNF protein within the spine head.

## 1. Introduction

Morphological changes in dendritic spines have consistently been observed during the process leading to the establishment of long-term synaptic changes and memory engrams (Zagrebelsky et al, 2020). Studies in the last two decades have largely clarified the mechanisms at the basis of these structural modifications. In particular, there is now evidence that the enlargement of the spine-head which accompanies synaptic long-term potentiation (LTP), a cellular correlate of memory formation, is entirely dependent on protein synthesis and the availability of neurotrophic factors such as the neurotrophin Brain-derived neurotrophic factor (BDNF) (reviewed in: Panja and Bramham, 2014; Leal et al., 2017; Zagrebelsky et al, 2020). Specifically, one early study showed that in order for LTP to be induced, BDNF must be locally released at synapses (Tanaka et al., 2008). A more recent study pointed out that when BDNF release occurs from post-synaptic terminals, LTP can be induced by activating an autocrine loop through TrkB receptors located on the very same post-synaptic spine compartment from which BDNF is released (Harward et al, 2016). Nevertheless, despite all these studies, the question of whether the post-synaptically released BDNF is locally synthesized in this compartment or has other origins, still remains unanswered (Leal et al, 2014; Song et al., 2017).

Studies in rodent hippocampal cultures have shown that BDNF is preferentially released from dendritic compartments in an activity-dependent manner upon Ca^2+^ entry through post-synaptic NMDA receptors (Kuczewski et al, 2009; Matsuda et al., 2009). Hence, it can be logically presumed that in response to LTP induction, BDNF is synthesized in dendritic spines or in the dendritic shaft near the activated synapses. However, there is also evidence that BDNF can be endocytosed at the postsynaptic level and further released after LTP induction (Santi et al., 2006; Wong et al., 2015). Surprisingly, studies using a genetically modified mouse in which endogenous BDNF was HA-tagged, reported the presence of this chimeric BDNF only in the pre-synaptic compartment (Matsumoto et al., 2008; Dieni et al. 2012, reviewed in Song et al., 2017). On the other hand, in support of the model posing that BDNF also has a post-synaptic localization, several research groups have gathered a large body of evidence at functional, immunofluorescence and electron microscopy level demonstrating BDNF localization within post-synaptic terminals (Edelman et al., 2015; Harward et al., 2016, Song et al., 2017; Brigadski and Leßmann, 2020).

One additional support to the view that BDNF localizes in post-synaptic terminals has come from our own and other groups studies showing that BDNF mRNA is present in dendrites, but not in axons, and its transport towards the distal dendritic compartment is enhanced in vivo in response to seizures, antidepressants, physical activity (Tongiorgi et al., 2004; Baj et al., 2012) and in vitro, in response to electrical activity, and treatment with NT3 or BDNF itself (Tongiorgi and Baj, 2008; Oe and Yoneda, 2010; Vicario et al, 2015; Mallei et al., 2015; Lekk et al., 2023). As a corollary to these findings on dendritic BDNF mRNA, chimeric BDNF-GFP constructs expressed in cultured neurons show local translation in dendrites in response to electrical activity (Baj et al., 2011; Vaghi et al., 2014 Lekk et al., 2023) and in experimental animals, endogenous BDNF protein and mRNA have an overlapping distribution in the hippocampal laminae containing the dendrites, both in resting conditions and following various types of stimuli (Tongiorgi et al., 2004; Baj et al., 2012). However, it still remains unclear whether the local translation of BDNF occurs in dendrites, or within the spine.

Several mRNA coding for various classes of proteins are known to be targeted to dendrites in mature neurons (Holt et al., 2019). Among them there are proteins of synaptic elements, translation factors, cytoskeleton, RNA-binding proteins and components of signalling pathways, including neurotrophic factors like BDNF (Holt et al., 2019; Glock et al., 2021). It has recently been shown that abundant dendritic mRNAs, such as the one encoding Calmodulin-Kinase II (CaMKII) or Rgs4, can move both forward and backward but, following stimulation of the cultures, they largely become stationary in dendrites and may eventually be locally translated in the vicinity of activated synapses (Bauer et l., 2019; Donlin-Asp et al., 2021). The question which has been put forward by an insightful review article by Rangaraju, Tom Dieck and Schuman (2017) is how strictly compartmentalized is the local production of a protein arising from translation of a dendritic mRNA. In other terms, is there a widespread translation of an mRNA, spanning a large segment of a dendrite, or is it a phenomenon restricted to the immediate neighbourhood of a spine?

In this study, we investigate these unresolved questions. Using chimeric constructs ectopically expressed in primary hippocampal cultures, we tracked BDNF mRNA trafficking within the dendrites and its local translation following induction of chemical-LTP (cLTP) in living mouse hippocampal neurons.

## 2. Materials and Methods

### 2.1 Primary hippocampal neuronal cultures

The animal study was approved by Organismo Preposto al Benessere degli Animali (OPBA) of the University of Trieste and by the Italian Ministery of Health. The study was conducted in accordance with the local legislation and institutional requirements. Dissociated hippocampal neurons were prepared from P0-P1 C57BL6/J mice pups. Hippocampi were dissected out in cold HBSS medium (Minimal Essential Medium, 0,2% MOPS, 20mM glucose, 1mM sodium pyruvate) and then incubated in 300μl of 0,25% Trypsin in HBSS at 37°C for 8 minutes. After 5 min of centrifugation at 8000 rpm, cells were mechanically dissociated in DMEM (DMEM 1x +5% of FBS) by pipetting up and down for 10 times with a Pasteur pipette. Cells were counted with the dye exclusion method using Trypan Blue (Sigma) in the Burker chamber (Eppendorf), and 400,000– 500,000 cells were obtained from each mouse. Cells were plated either on 12 mm cover slides coated with 0.1mg/ml poly-L-ornithine(Sigma), or on µ-Slide 8 Well Glass Bottom (#1.5 polymer coating cat. No. 80806 from IBIDI) at a density of 130,000 cells/well. In order to reach this density, having different coverslips per experiment, a pool of hippocampi from different animals were used for each preparation. After one hour from plating, DMEM was replaced with Neurobasal medium supplemented with 2% B-27 (Invitrogen), 1 mM Ala-Gn (Glutamax, G1845, Sigma) and 0.45% Glucose and 1% penicillin-streptomycin. A final concentration of 1.25 uM Ara-C was added at days-in-vitro 4 (DIV4) to inhibit glia proliferation. Neurons were cultured under a humidified 5% CO_2_ atmosphere at 37°C for 14 days. At DIV4 half medium was replaced by adding Ara-C. Neurons were transfected in 8 µ-Slide 8 Well Glass Bottom chamber slides (IBIDI) with 1.2 µl of DNA and 1.2 µl Lipofectamine 2000 per well and 50 µl of MEM. To study BDNF mRNA trafficking, neurons were transfected between DIV11-13 with EX6-BDNFcds-12xMS2-3L and MS2-NLS-mcherry or Ex1-BDNFcfds-12xMS2-3L and MS2-NLS-mcherry or with CaMKIIa-12xMS2-3’UTR together with MS2-NLS-mcherry. To study the BDNF protein, transfections of EX6-BDNFcds-gfp-3L or Ex1-BDNFcds-gfp-3L were carried out at DIV9 and cultures were visualized at DIV13-14.

### 2.2. cLTP protocols and solutions

Long-term potentiation was chemically induced by incubation for 30, 60 and 90 minutes with 10 μM Forskolin (Sigma 344270) in a solution containing 92.4 mM NaCl; 2.3 mM KCl, 1.3 mM CaCl_2,_ 0.35 mM Na_2_HP0_4,_ 4.2 mM NaHCO_3,_ 0.45 mM K_2_HPO_4,_ 7 mM HEPES, 5.5 mM D-glucose. The control solution contained 92.4 mM NaCl; 2.3 mM KCl, 0.8 mM MgSO_4,_ 1.3 mM CaCl_2,_ 0.35 mM Na_2_HP0_4,_ 4.2 mM NaHCO_3,_ 0.45 mM K_2_HPO_4,_ 7 mM HEPES, 5.5 mM D-glucose. The pH was 7.4. Solutions were pre-warmed before the experiments.

### 2.3. Live imaging protocol for the BDNF mRNA granules

Neurons plated in IBIDI chamber slides were transfected as described at point 2.1 with EX6-BDNFcds-12xMS2-3L and MS2-NLS-mcherry or Ex1-BDNFcfds-12xMS2-3L and MS2-NLS-mcherry and time-lapse video images were taken 17-18 hours after transfection. To acquire movies, a Nikon Eclipse Ti-E-epifluorescence microscope using 40Xobjective (1.0 NA oil PlanApo DICH) equipped with a DS-Qi2 Camera (NIKON) was used. Movies of 10 minutes at 1 frame/second for control condition and of 15 minutes at 1 frame/second for the cLTP condition were acquired. During the 10 minutes of control recordings, neurons were in the control medium (as described at point 2.2). Then, a wash with the medium without MgSO4 was done. For stimulating neurons, 10μM of forskolin was added to the medium without MgSO4 and neurons were imaged for further 15 minutes.

### 2.4. BDNF protein transfection and visualization in live

Neurons coated in IBIDI chamber slides were transfected with ex1-BDNFcds-GFP-3’UTR long or ex6-BDNFcds-GFP-3’UTR long or BDNFcds-GFP. In order to visualize the dendrites and the dendritic spines, neurons were co-transfected with an mCherry filler. Video time-lapse images were taken at 17-18 hours from transfection. Nikon Eclipse Ti-E-epifluorescence using 40X objective (1.0 NA oil PlanApo DICH) equipped with a DS-Qi2 Camera (NIKON) was used to acquire the movies. Recordings of 10 minutes at 1 frame/minute for control condition and of 60 minutes at 1 frame/minute for activated condition were acquired. During the 10 minutes of control condition, neurons were in the control medium (as described at point 2.2.). Then a wash with the medium without MgSO4 was done. For stimulating neurons, 10μM of forskolin was added to the medium without MgSO4 and neurons were imaged for further 60 minutes.

### 2.5. Video time-lapse analysis for the BDNF mRNA and protein

After the acquisition, movies were opened on ImageJ and Kymographs were extracted using the plugin Multi Kymograph. Once the Kymographs were extracted, the analysis consisted in selecting a particular movement of the granules by drawing a segmented line over the corresponding track of the kymograph. Then, thanks to a custom ImageJ MACRO the position and the time coordinates (x and t) from the segmented line were extracted. Thus, all the necessary information about the dynamics of the granules, as the average and instantaneous velocities and the overall behaviour of the particle (anterograde, retrograde or confined) were extracted. The net movement of a granule was considered. Confined movements were defined for granules that moved less than 0.5 μm. For the video of the BDNF protein only neurons transfected with BDNFcds-GFP were then analysed because with other constructs neurons were not well visible. The mean fluorescence of BDNFcds-GFP protein spots was measured every 10 minutes during the 60 minutes of Forskolin treatment.

### 2.6. Evaluating the distance between a granule and a synapse

After 17-18 hours from transfection with BDNF mRNA, neurons were stimulated with 10 μM Forskolin for 15, 30 and 60 minutes. Then neurons were fixed and labelled with anti-Synapsin I antibody (Rabbit, dil. 1:1000, Millipore cat. no. AB1243), and revealed by anti-rabbit secondaru antibodies (Alexa 488 made in goat, dil. 1:500; ThermoFischer Scientific, cat. No. A11034). Images were acquired using Elyra 7 microscope using 63X objective using a filter combination: BP 420-480 and BF 405/488/561/642. SIM images were reconstructed through the Zen-black software. A SIM reconstruction followed by a Z-stack projection was performed using Zen-black software. A threshold for the channel of Synapsin I was applied and a mask was created. The same was done for the BDNF mRNA channel. In this latter case, two thresholds were considered: one threshold for identifying the granules and a second threshold for delineating the structure of the neuron with the relative spines. A MACRO for speeding up the analysis was made. Once the thresholded and masked images were created, stretches of neurons between 30-50 μm with visible spines were considered. Granules were counted between 3 μm (away from the soma) and -3μm (closer to the soma) from an longitudinal axes that divides in two parts the spine. So, all granules within this interval of a total of 6 μm were considered. Granules inside the spines were also counted. Stretches of 30-50 μm were considered of apical distal dendrites.

### 2.7. Statistical analysis

Data analysis was performed blind for all experiments. Statistical values are represented as mean ± S.E.M when data are parametric. When data are not parametric, median ± 95% confidence was chosen. The number of experiments and cells analysed for each situation is indicated in each figure. Statistical significance was calculated using GraphPad Prism version 8 (GraphPad Software, LaJolla, USA). Normality distributions test was done by Shapiro-Wilks test. If samples proved to have a normal distribution, then Student t-test (for two groups) or ANOVA with Tukey’s post-test (for three or more groups) were used. When samples did not have a parametrical distribution, then for experimental conditions with two data sets the Mann-Whitney test was used or the Krustal-Wallis for three or more groups. χ² statistical analysis was also performed in order to identify independence between categories. Cumulative distribution of data was also used. Outliers were removed using the formula with the interquartile range rule. This required to calculate the first quartile *Q*_1_ and the third quartile *Q*_3_ and the difference between them. The difference was then multiplied by 1.5. It was necessary to add 1.5x (*Q*_3-_ *Q*_1_) to the third quartile and to subtract 1.5x (*Q*_3-_ *Q*_1_) any number greater or smaller than these values was a suspected outlier. Significance was set as *p<0.05; **p<0.01; ***p<0,001. The p value is reported in each figure legend in the result section, as the statistics used for each experiment.

## 3. Results

### 3.1. Time-course for the visualization of exogenous mRNA after transfection

The MS2 loop system is a well-known method for imaging mRNAs trafficking in living neurons (Dahm et al., 2008). It consists in the insertion within a gene of interest of a repeat of several identical stem-loop structures derived from the bacteriophage RNA, associated with the ectopic expression of fluorescently-tagged MS2-coat binding protein (MCP) which binds to the MS2 loops. Since previous studies reported that overexpression of MCP is potentially prone to artifacts (Dahm et al., 2008; Yoon et al., 2016; Bauer et al., 2019), in a first set of experiments we carried out a time course analysis to find the optimal time point to image neurons and avoid possible artifacts. For this purpose, neurons were co-transfected (DIV 11-12) with the plasmid encoding the positive control gene CamkIIα bearing eight MS2-loops in its 3’untranslated region (CamKII-8L-3’UTR) and a second plasmid encoding the MS2-Coat Protein (MCP) in fusion with the fluorescent reported gene mCherry (MCP-mCherry). As a negative control, neurons were transfected with only the plasmid encoding MCP-mCherry. Neurons were imaged at different time points from transfection, i.e. 10-12, 17-18, 22-24, 36, or 68 hours (h). In all conditions, 10 minutes-long videos were recorded for each neuron, and the movements of fluorescent mRNA granules analysed. In control experiments, MCP-mCherry granules were detected starting from 22-24h and were not visible at earlier times (MCP_alone, **Figure 1C**). The number of neurons with MCP-mCherry granules increased significantly over time (**Figure 1C**): 15% of neurons showed granules at 22-24h (## p = 0.0015 vs. 10-12h), 40% of neurons after 36h (### p = 0.0002 vs. 10-12h), and 100% after 68h from transfection (#### p ≤ 0.0001 vs. 10-12h, n= 3 independent cultures). The number of granules detected in each neuron (**Figure 1D**) also increased over time, from an average of 1.68 granules/neuron (22-24h), to 2.83 granules/neuron (36h), and 4.18 granules/neuron (68h). The ANOVA test followed by Tukey’s multiple comparison indicated that this increase was significant only at later stages (22-24h vs. 36h p = 0.0910, ns; 22-24h vs. 68h p = 0.0002 ***; 36h vs. 68h p = 0.0056 **). Of note, the diameter of the granules, taken as a measure of granule size, remained unchanged over time (**Figure 1E**; 22-24h vs. 36h p= 0.89; 22-24h vs. 68h p=0.88; 36h vs. 68h p=0.64).

**Figure 1.**
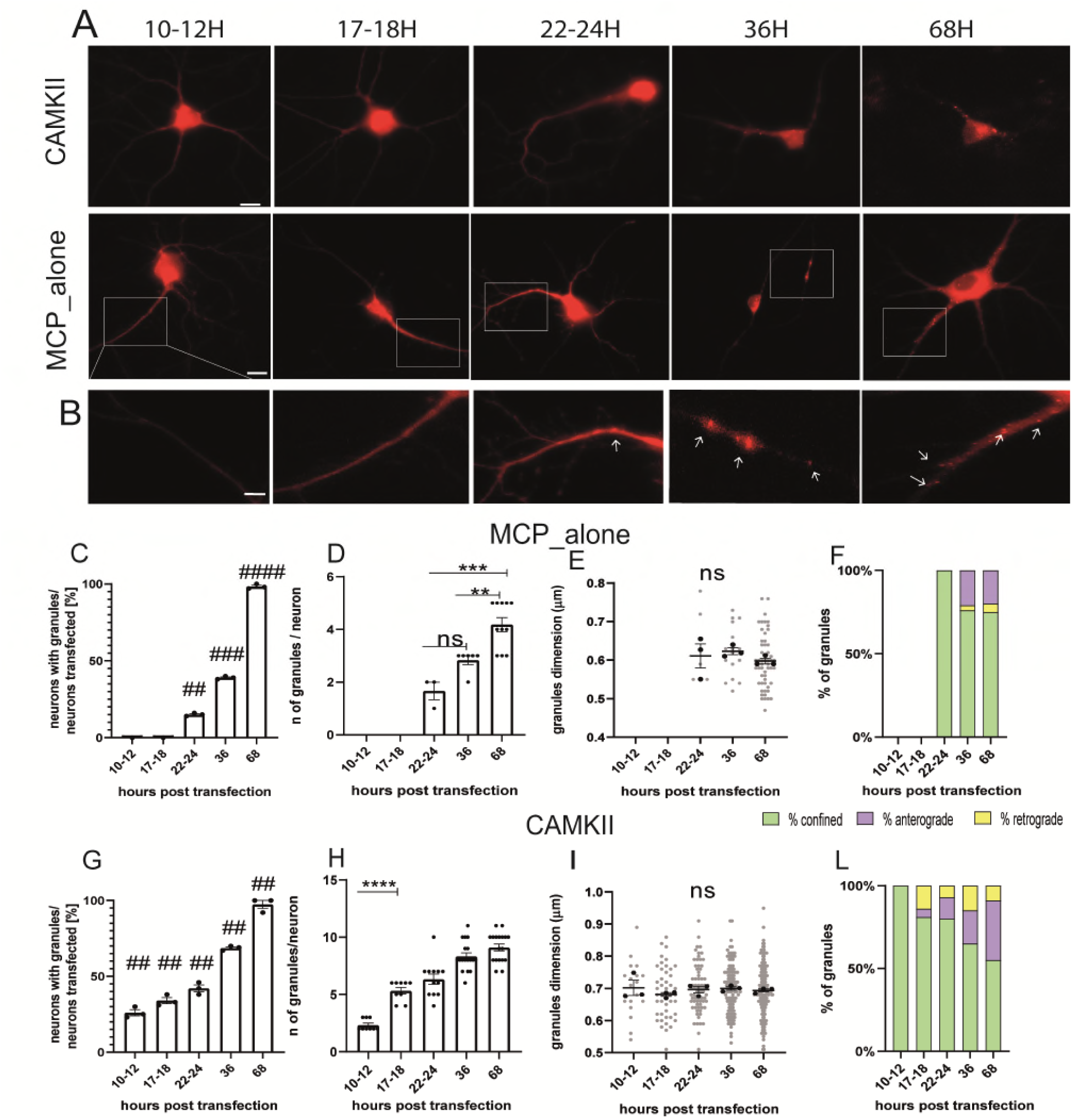
Settings for live-imaging analysis. A) Representative images of MCP_alone and CamkIIα in hippocampal neurons at different times post transfection B) High-magnification images of neurons transfected with MCP_alone. Arrows indicate granules. C) Number of neurons with granules after different times post transfection (t test); D) Number of granules present in the neurons transfected after different times. Note that before 22-24 hours granules were not detected (ANOVA followed by Tukey’s post hoc test). E) Dimension of the MCP_granules (ANOVA followed by Tukey’s post hoc test). F) Movement dynamics of granules (χ² test). G) Number of neurons with granules after different times post transfection (t test). H) Number of granules present in the neurons at the different time points from transfection. I) Dimension of CamkIIα granules (ANOVA followed by Tukey’s post hoc test). L) CamkIIα movements dynamics after different hours post transfection (χ² test; p<0.001 at 10-12 and 17-18-hours post transfection). For each graph n=15-45 neurons from 3 independent experiments were analysed.

Live imaging analysis to evaluate the dynamics of MCP-mCherry mRNA granules was carried out using ImageJ (Multi Kymograph plugin) and, depending on type of movement observed, each granule was assigned to the class of either anterograde granules (moving prevalently away from the cell body), retrograde granules (moving towards the cell body) or confined granules (with movements with total displacement of less than 0.5 μm). Live-cell videos were acquired at 1 frame/sec using a 40X oil objective. Dynamic analysis showed a differential granule behaviour (confined, anterograde and retrograde) at different time points (χ² test p<0.0001) (**Figure 1F**). Specifically, at 22-24h, 100% of granules were confined; at 36h granules were 76% confined, 21% anterograde, 3% retrograde, and at 68h were 75% confined, 20% anterograde and 5% retrograde. For each plot: n=15-45 neurons from 3 independent cultures (5-15 neurons, each). Taken together, these data indicate that if the imaging of neurons is conducted before 22-24h from transfection, the granules produced by the aggregation of the MCP protein alone are not yet formed, thus suggesting that 10-12h or 17-18h post-transfection were the preferable time windows to image neurons in order to avoid this artefact.

The same analysis was performed also for neurons co-transfected with the CamKII-8L-3’UTR and the MCP-mCherry plasmids (**Figure 1** **G**). Granules were visible in 26% of neurons already after 10-12h, in 34% of neurons after 17-18h, in 42% after 22-24h, in 68% after 36h and in 97% after 68h (Tukey’s multiple comparison test Significance Adjusted p value 10-12h vs. 17-18h ns p>0.9999; 10-12h vs. 22-24h ***p=0.0002; 10-12h vs. 36h ****p≤0.001). The number of CamKII-8L-3’UTR granules detected (**Figure 1H**) increased from an average of 2.33 granules/neuron at 10-12h to 5.3 granules/neuron at 17-18h, 6.3 granules/neuron at 22-24h, and 8.3 granules/neuron at 36h, reaching 9.11 granules/neuron at 68h. CamKII, granules diameter remained unchanged over time (**Figure 1I**, n= 3 independent cultures, n=5-15 neurons for each culture). Considering the dynamics of the granules, the large majority of them resulted confined. Specifically, at 10-12h, 100% granules were confined; at 17-18h granules were 81% confined, 5% anterograde, and 14% retrograde; at 22-24h, 80% were confined, 13% anterograde, and 7% retrograde; at 36h, 65% of granules were confined, 20% anterograde and 25% retrograde; and at 68h, were 55% confined, 36% anterograde and 9% retrograde (**Figure 1L**). The χ² test showed a significantly different granule motility between 10-12h and 17-18h (p<0.0001). Considering these results, the optimal time point to image neurons resulted to be 17-18 hours post-transfection, since the number of CamkIIα granules was sufficiently large (5 granules/neuron) as compared to 10-12h (2 granules/neuron, 10-12h vs 17-18h, p<0.0001), and without artefacts for the MCP-mCherry alone. Moreover, the live imaging experiments indicated the 17-18h time point as suitable for dynamic analysis because granules were moving in both anterograde and retrograde direction.

### 3.2. Tracking of BDNF mRNA trafficking in living neurons

Once the settings for visualizing exogenous mRNA were established, hippocampal neurons were transfected with BDNF mRNA constructs and the MCP-mCherry at 11 DIV and living neurons were imaged on the following day i.e., at 17-18 hours post-transfection. The very same neuron was first imaged in control solution (see Materials and Methods) for 10 minutes (min) to establish the baseline and then, for 15 min in a solution containing 10 µM forskolin. To assess BDNF mRNA dynamics, we focused on the BDNF transcript which has the most prominent dendritic localization, i.e. with the 5’UTR encoded by BDNF exon 6 and the long version of the 3’UTR, and we compared it with the most somatically restricted BDNF transcript, i.e. the one with the 5’UTR encoded by BDNF exon 1, also having the long 3’UTR (Chiaruttini et al., 2009; Vicario et al., 2016; Colliva et al., 2022). In both BDNF transcripts, a 12 MS2 loops repeat was inserted at the end of the coding sequence (CDS) generating the constructs Ex6-bdnfCDS12L-3UTR, and Ex1-bdnfCDS-12L-3UTR. As positive control, the CamkIIα-8L-3UTR construct was used, and as negative control a plasmid carrying only the 12 MS2 stem loop (12L) was generated. This last plasmid does not have neither a coding sequence, nor a 3’UTR sequence, but just the same bacteriophages stem loops which were cloned in the BDNF and in CamkIIα constructs.

The dynamics of CamkIIα-8L-3UTR, Ex6-bdnfCDS12L-3UTR, Ex1-bdnfCDS-12L-3UTR, and 12L mRNA trafficking was evaluated (**Figure 2A**). Granules, whose typical appearance is shown for Ex6-bdnfCDS12L-3UTR in **Figure 2B**, were found in both apical and basal dendrites (**Figure 2C**) but with some differences in the percentage of localization between these two compartments, depending on the mRNA analysed. Specifically, 48% granules showed an apical localization and 52% basal for CamKII, 56% apical and 44% basal for Ex6, 66% apical and 34% basal for Ex1 and 71% apical and 29% basal for the 12L.

**Figure 2.**
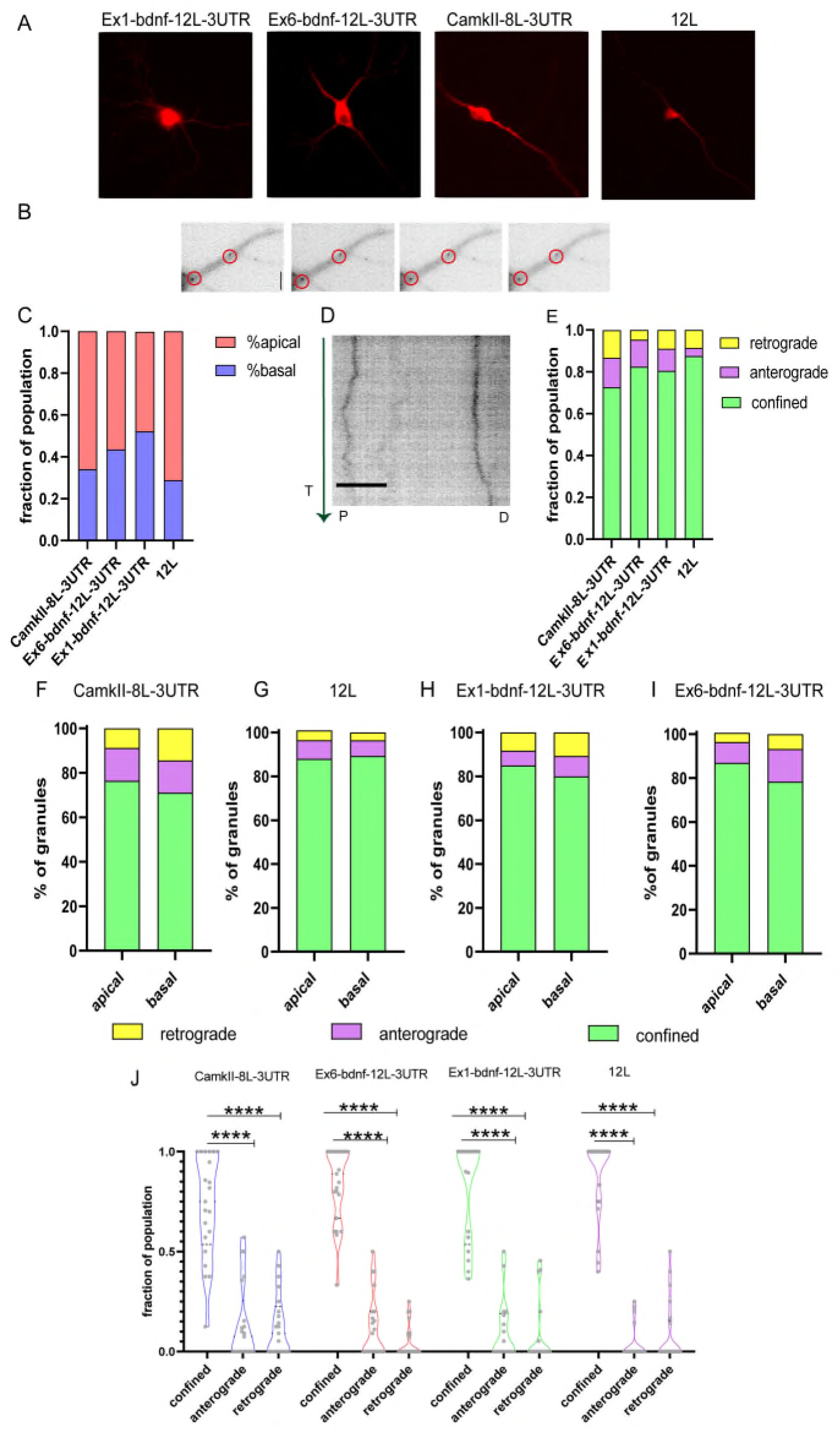
Tracking mRNA granules in live neurons. A) Representative neurons transfected with the BDNF mRNA construct and MCP_mcherry, or CamkIIα and MCP_mcherry or the 12L construct. B) Representative dendrites with Ex6-bdnf-12L-3UTR granules encircled. C) Fraction of population of granules present at apical and basal dendrites. D) Representative kymograph of a granule movement: the green vertical line represents the time (T) and the arrow indicate the time direction from the beginning of the analysis. P and D represent respectively the Proximal and Distal part of the dendrite. E) Fraction of granules population showing the movement dynamic: the vast majority of granules show confined behaviour (ANOVA followed by Kruskal Wallis post hoc test). F) Apical and basal granules of CamkIIα show similar distribution of anterograde, retrograde and confined granules behaviour without any statically difference by the χ² test, the same test was applied for G) 12L apical and basal granules and H) Ex1-bdnf-12L-3 UTR and I) Ex6-bdnf-12L-3UTR showing that apical and basal granules for each construct behave similarly without any significant statistical change. J) Confined, anterograde and retrograde fraction of population. The vast majority of granules were confined (ANOVA followed by Kruskal Wallis post hoc test) did notshow significant difference between the different constructs. Data shown were acquired from three independent experiments and a total of 20 neurons per condition were imaged.

From the analysis of the kymographs obtained from apical and basal dendrites (**Figure 2D**) it resulted that the large majority of the granules formed by the four constructs were confined, with the control construct 12L having the highest percentage of confined granules (**Figure 2E**), in analogy to previous reports (Doyle and Kiebler, 2011; Yoon et al., 2016). The graph in **Figure 2E** shows the fraction of granules displaying confined, anterograde, retrograde movements within the total granule population for each analysed neuron. Regarding the active movements, the three constructs, with the exception of 12L, displayed a slight bias, not statistically significant, for anterograde transport (fraction of population, anterograde vs. retrograde: CamKII: 0.075 upper limit 0.57 and lower limit 0 vs 0.09 upper limit 0.5 and lower limit 0; Ex1-bdnf-12L-3UTR: 0 upper limit 0.5 and lower limit 0 vs 0 upper limit 0.25 and lower limit 0; Ex6-bdnf-12L-3UTR: 0.5 upper limit 0 lower limit 0 vs 0.45 upper limit 0 lower limit 0). For the construct 12L, the anterograde and the retrograde transport showed similar results (median:0 upper limit 0.25 and lower limit 0 for the anterograde; 0 upper limit 0.5 and lower limit 0 for the retrograde). A statistical comparison for each granule classes across the different constructs suggested no significant differences, except for a non statistically significant trend for a higher confined population for the 12L granules with respect to CamkIIα (**Figure 2E**; n= 20-21 neurons for each construct, p=0.13). Next, we verified if the type of granules movement was influenced by their localization in the basal or apical dendrites (**Figure 2F-I**). The χ² statistical test indicated that the percentage of movements showed by the RNA granules was not significantly different between apical and basal dendrites, for all constructs (**Figure 2F-I**). Since basal and apical dendrites showed similar percentages of confined, anterograde and retrograde granules, in all subsequent analyses the data from the two compartments were pooled together (**Figure 2J**), showing that the majority of granules were confined.

The distribution of the granules within the dendrites was also evaluated by counting the number of granules found in dendritic segments at increasing distance from the soma, namely at <20 µm, between 20-6 µm, or 60-100 µm, and >100 µm from the soma (**Figure 3A**). Histograms of granules distribution in neurons in resting conditions, clearly show that the majority of granules for the four constructs was localized in the proximal dendrites, i.e. between 20-60 µm from the cell soma (**Figure 3B-F**).

**Figure 3.**
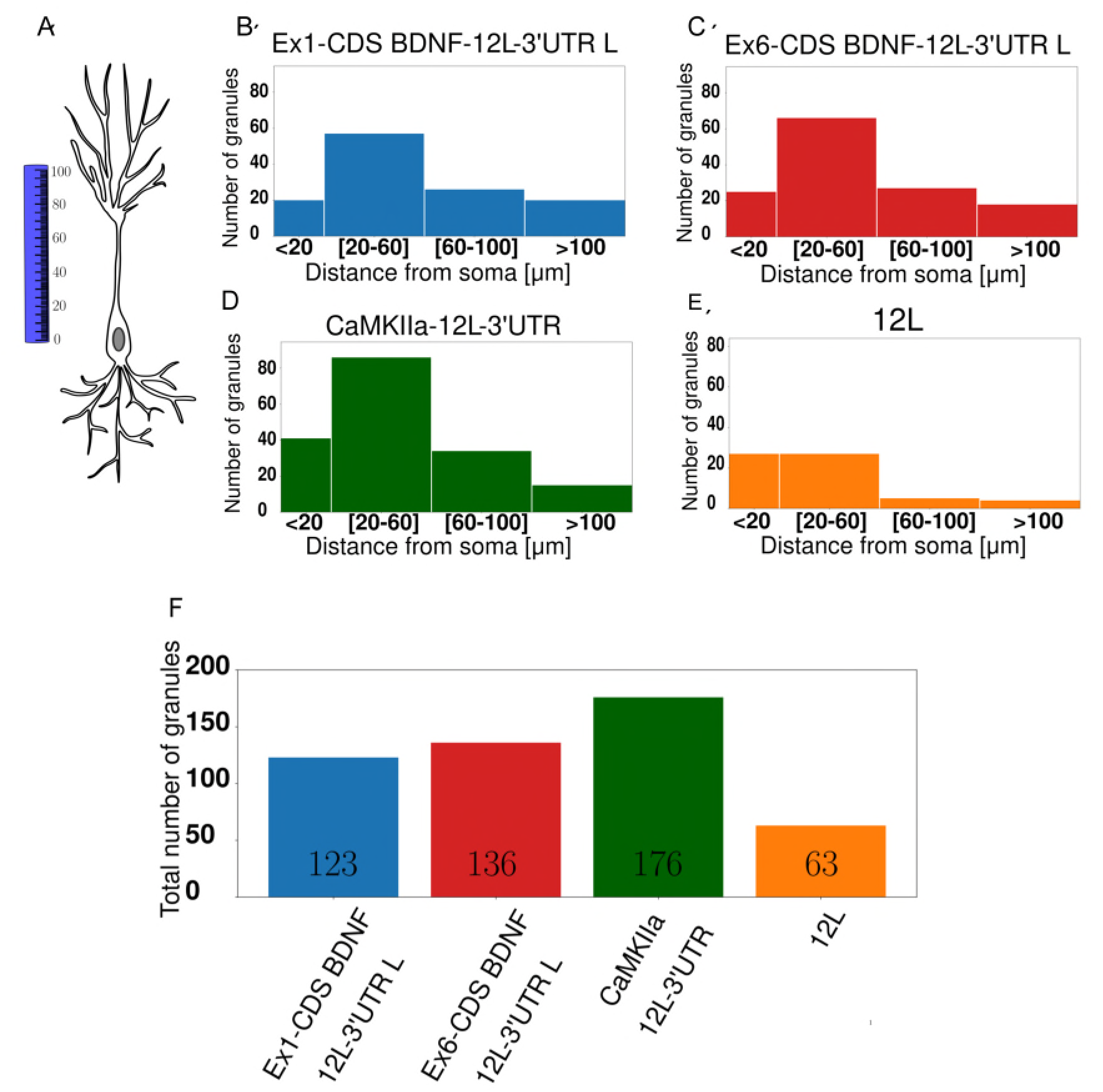
Subcellular localization of mRNA granules. A) Graphical representation showing how the distance from the soma was calculated B) Ex1-cdsBDNF-12L-3UTR histogram with the number of granules detected at proximal (20-60 μm) distal (60-100 μm) dendrites, the majority of granules is at 20-60 μm C) Ex6-cdsBDNF-12L-3UTR histogram D) CamkIIα-12L-3UTR histogram even in this case the majority of granules is at proximal dendrites E) 12L histogram F) Total number of granules detected in almost 20 neurons for each construct.

### 3.3. Analysis of granule movements in resting neurons

The population of granules with anterograde or retrograde active movements was analysed for their velocity and the distance travelled during the observation time. The four constructs showed similar velocities (**Figure 4A-C**). For the anterograde movements (**Figure 4B**), velocities were: CamKII: 0.153 µm/sec, Ex1-bdnf-12L-3UTR: 0.133 µm/sec, Ex6-bdnf-12L-3UTR: 0.189 µm/sec, 12L: 0.125µm/sec. For the retrograde movements, velocities were: CamKII: 0.234 µm/sec, Ex1-bdnf-12L-3UTR: 0.195 µm/sec, Ex6-bdnf-12L-3UTR: 0.143 µm/sec, 12L: 0.176 µm/sec (**Figure 4C**). The observed velocities are consistent with similar values previously reported in the literature (Chiaruttini et al., 2009a) and the only statistically significant difference was found between Ex6-bdnf-12L-3UTR and Ex1-bdnf-12L-3UTR (p =0.035) with the Ex6 construct showing a higher anterograde velocity than the Ex1 construct (**Figure 4B**), whereas for the retrograde transport there were no significant differences among the different constructs (**Figure 4C**). Also the travelled distance during the observation time was similar for all mRNA constructs (**Figure 4D**). For the anterograde movements the travelled distance was CamKII = 3.5 µm, Ex1-bdnf-12L-3UTR = 3.9 µm, Ex6-bdnf-12L3UTR = 3.4 µm, 12L = 5.3 µm (**Figure 4E**), while for the retrograde movement CamKII = - 10.790 µm, Ex1-bdnf-12L-3UTR = - 2.9 µm, Ex6-bdnf-12L3UTR = -3.1µm, 12L = -2.8 µm (**Figure 4F**).

**Figure 4.**
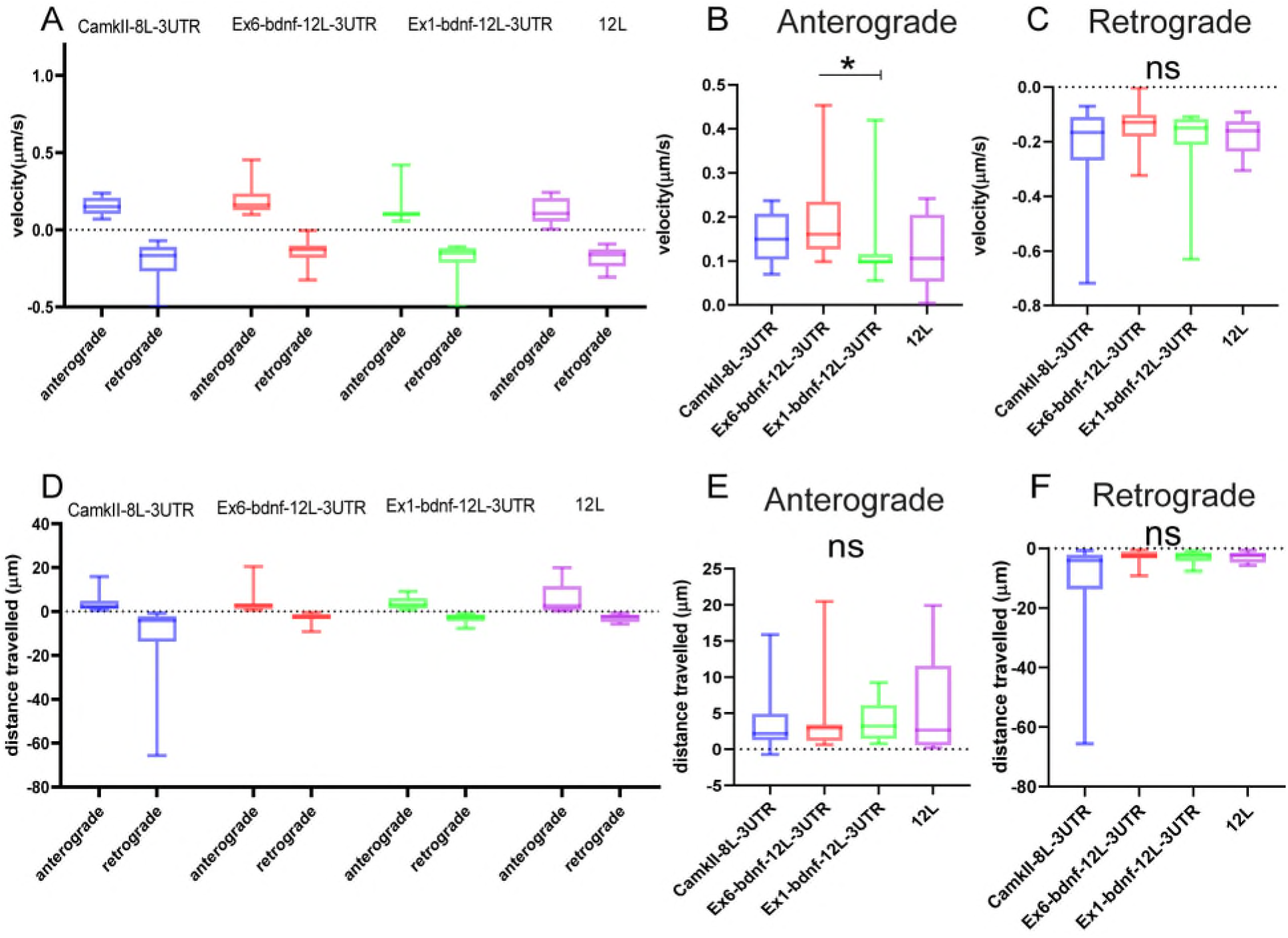
Type of movements of mRNA granules. A), B), C) Anterograde and retrograde velocity for the four constructs B) Significant difference between Ex6 and Ex1 for anterograde velocity (P=0.035) ANOVA followed by Kruskal Wallis. D) E) F) Total displacement of anterograde and retrograde movements. Data are represented by the median with 95% CI.

### 3.4. Analysis of granule movements after cLTP induction

After 10 minutes of baseline imaging, 10 µM forskolin was added to chemically induce LTP (cLTP), and each neuron was imaged for an additional 15 minutes (**Figure 5A**). Following cLTP induction, granules exhibited a movement arrest and showed confined behaviour (**Figure 5B, C, D, E**; n= 20-21 neurons). A paired t test statistical analysis showed that, comparing the behaviour of the same granules in the ctrl and cLTP condition, the confined population was significantly increased after cLTP for Ex6 (p= 0.0234, panel 5B); ex1 (p=0.0156, panel 5C); CamkIIα (p=0.0084, panel 5D); and 12L (p=0.0156, panel 5E). For the anterograde population there was a significant decrease after cLTP with respect to ctrl, for Ex6 (p=0.0156, panel B), and Ex1 (p=0.0313, panel 5C), but not for CamkIIα (p=0.11, panel 5D), nor for 12L (p=0.12, panel 5E). After the cLTP induction, the retrograde population was significantly reduced for CamkIIα (p=0.0020, panel 5D) and 12L (p=0.0313, panel 5E). The finding that 12L construct exhibited a behaviour similar to that of the CamkIIα mRNA was surprising because the 12L construct was not expected to have a biological significance.

**Figure 5.**
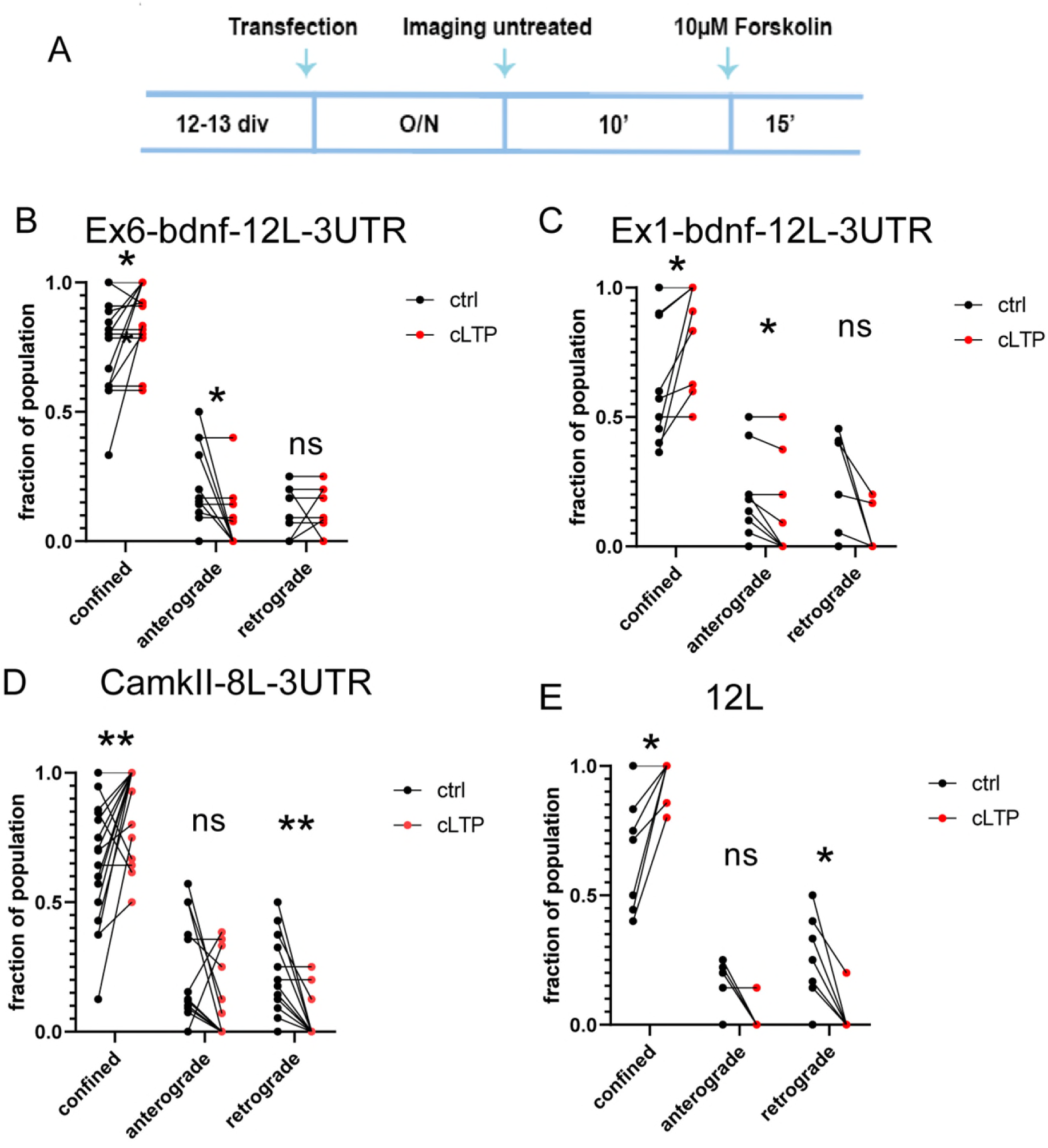
Dynamics of mRNA granules after cLTP induction. A) Experimental outline: after an overnight (O/N = 17-18h) from transfection, neurons were imaged for 10’ in control medium (untreated) and then, the same neurons were incubated with 10 μM Forskolin and imaged for additional 15’. B) After cLTP, Ex6-bdnf-12L-3UTR granules showed an increased fraction of confined granules (p=0.0234) and a decrease in the fraction of anterograde granules (p=0.0156). C) After cLTP, Ex1-bdnf-12L-3UTR showed an increase of the confined granule population (p=0.0156) and a decrease in the anterograde granules (p=0.0313). D) CamkIIα-8L-3UTR showed an increase in the confined granules (p=0.0084) and a decrease in the retrograde granules (p=0.0020). E) 12L showed an increase of confined granule population (p=0.0313) and a decrease in the retrograde granules (p=0.12) Statistical analysis was performed by paired t test.

### 3.5. cLTP-induced confinment of BDNF mRNA in proximity to a spine

From the live-cell analysis we concluded that after cLTP stimulation, mRNA granules arrested their anterograde movements becoming stationary. However, because of resolution limitations, it was not clear if after the cLTP-induced stop of mRNA trafficking, granules were accumulated in the proximity of a spine or not. To clarify this point, neurons transfected with the BDNF constructs Ex6-bdnfCDS-12L-3UTR, Ex1-bdnfCDS-12L3UTR and the control constructs CamkIIα-8L-3’UTR and 12L were analyzed with the super-resolution Elyra7 SIM microscope. Neurons were fixed at different time points, i.e. at 15 minutes, 30 minutes and 60 minutes from cLTP induction. Then, neurons were immunolabelled with Synapsin 1 to identify the position of pre-synaptic terminals and images were acquired (see Materials and methods). During post-processing analysis, thresholded images of the mRNA granules and Synapsin 1 were created (**Figure 6A**). Spines were identified as Synapsin 1 labeled protrusions along the dendrites. To identify mushroom and stubby spines, the head diameter and the neck of each spine were measured. Both spine types were considered for the analysis. For each dendrite, a ROI of 30-50 µm was drawn and all the spines in it were considered. The density of granules was counted in each ROI (granules/µm = number of granules per length unit), and the number of granules located within +3 µm (granules more distant to the soma) and -3 µm (granules closer to the soma) from each spine was considered. For this purpose, an orthogonal axis dividing the spine in two halves was designed as a reference line (see **Figure 6A**). Granules were also classified as being inside the spine head, or in the dendritic shaft segment below the spine. In both cases (inside or below the spine) a position closer to 0 indicates that the granule is positioned close to the spine axis which is generally running in correspondence to the spine neck (see **Figure 6A**).

**Figure 6.**
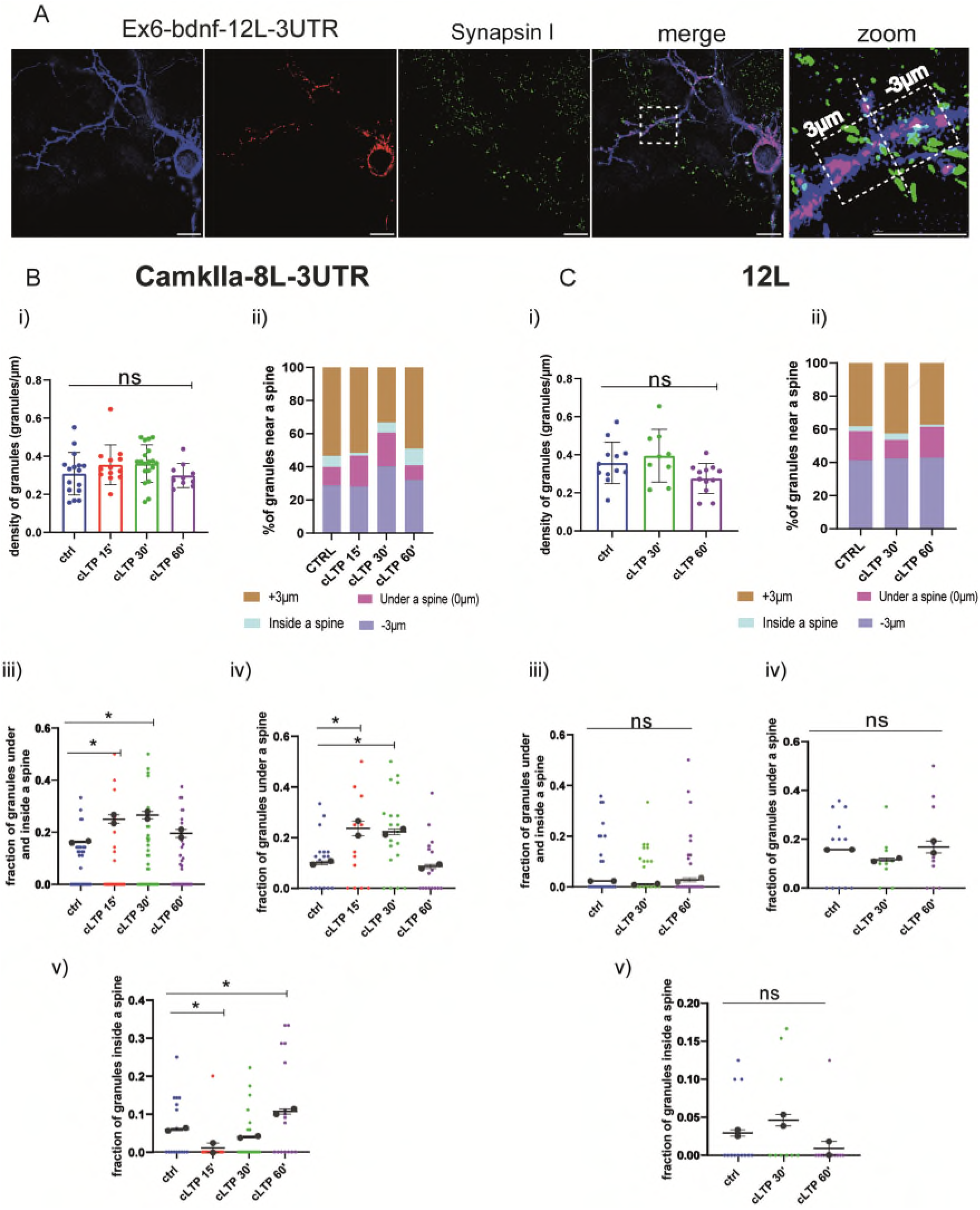
CamkIIα and 12L mRNA granules dynamics in proximity to dendritic spines. A) Representative thresholded images of Ex6-BDNF-12L-3UTR and Synapsin I. From the high magnification section a spine is visible, granules found in the range of +3 μm and -3 μm were considered. Scale bar 5 μm. **B) CamkIIα-8L-3UTR granules** i) CamkIIα mRNA granules counted in each dendrite streches of 30-50 μm. ANOVA followed by Dunnett’s post hoc test revealed no significant difference in the density of mRNA CamkIIα granules after cLTP stimulation ii) mRNA granules distribution near a spine after cLTP induction. Granules under a spine and inside a spine were considered iii) CamkIIα granules under and inside a spine were considered at different cLTP induction. After 15’ and 30’ from cLTP induction there is an increase in the granules present under and inside a spine (ctrl vs cLTP 15’ p=0.0311, ctrl vs cLTP 30’ p=0.0180) iv) CamkIIα granules under a spine were significantly higher after 15’ and 30’ from cLTP induction (p= ctrl vs cLTP 15’ p=0.0140 and ctrl vs cLTP 30’ p=0.0204) v) CamkIIα granules inside a spine were significantly higher after 60’ from cLTP ctrl vs 15’ p=0.03369 and ctrl vs cLTP 60’ p= 0.00336) **C) 12L granules** i) 12L mRNA granules counted in each dendrite streches of 30-50 μm ANOVA followed by Dunnett’s post hoc test revealed no significant difference in the density of 12L granules after cLTP stimulation ii) mRNA granules distribution near a spine after cLTP induction iii) 12L granules under and inside a spine were considered at different cLTP induction. iv) 12L granules under a spine were not significantly higher after cLTP induction v) 12L granules inside a spine were not significantly higher after cLTP.

The density of granules for CamkIIα and 12L counted in each dendrite stretches of 30-50 μm, did not change at the different time points analysed after cLTP induction (Panel (i) in **Figure 6B** and **6C** for CamkIIα and 12L, respectively). However, when considering the CamkIIα granules located under the spine and those found inside the spine as a single fraction of the whole population, a significant increase was found at 15’ and 30’ after cLTP induction, with respect to baseline (**Figure 6B, C** Panel iii, ctrl vs cLTP 15’ p=0.0311, ctrl vs cLTP 30’ p=0.0180; summarized as % in Panel ii). Interestingly, when the two fractions of CamkIIα granules located under and inside the spine were considered separately, an opposite behaviour was found. In detail, the fraction of granules under the spine transiently increased at 15’ and 30’ to return to baseline levels at 60’ after cLTP (**Figure 6B**, Panel iv, ctrl vs cLTP 15’ p=0.0140, and ctrl vs cLTP 30’ p=0.0204;.Summarized as % in Panel ii), while the fraction of CamkIIα granules located inside a spine was significantly decreased in the early phase after cLTP (15’), to become significantly higher than the baseline in the late phase, at 60’ post-cLTP (**Figure 6B**, Panel v, ctrl vs 15’ p=0.03369 and ctrl vs cLTP 60’ p= 0.00336; summarized as % in Panel ii). As expected, the negative control 12L did not distribute differently inside and under a spine in control and after cLTP stimulation granules (**Figure 6C**, Panels i-v).

Considering the BDNF constructs, also for Ex1-bdnfCDS-12L-3UTR (BDNF-Ex1) and Ex6-bdnfCDS-12L-3UTR (BDNF-Ex6) there was no statistically significant change, with respect to baseline, in the granule’s density at the various time points after cLTP (**Figure 7A** for BDNF-Ex1, and **7B** for BDNF-Ex6, Panels i and ii). A significant increment in the pooled fraction of granules inside and under a spine was observed after 30’ from cLTP induction (ctrl vs cLTP 30’ p=0.0266), but not at 15’ or 60’ (ctrl vs 15’ p=0.22, and ctrl vs 60’ p=0.13). When considering the granules inside and under the spine separately, there was an increase in BDNF-Ex1 mRNA granules under a spine after 15’ from cLTP (**Figure 7A**, panel iv; ctrl vs cLTP 15’ p=0.0047), and an increase in BDNF-Ex1 mRNA granules inside the spine after 30’ from cLTP (ctrl vs 30’ p=0.0253; **Figure 7A**, panel v). For the BDNF-Ex6 construct, the pooled fraction of granules inside and below a spine, was significantly higher at 15’ with respect to 60’ (p=0.0115), but not with respect to the control condition (p=0.0785; **Figure 7** **B,** panel iii). Looking at these granule populations separately, there was an increase in the fraction of granules under a spine after 15’ from cLTP induction (**Figure 7** **B**, panel iv; ctrl vs 15’ cLTP p=0.0005), while the fraction of granules inside the spine was unchanged (**Figure 7** **B**, panel v, ctrl vs cLTP 60’ p=0.1271). Taken together, these results suggest that the distribution of BDNF mRNA within the dendrites changes after cLTP induction with a significant accumulation of BDNF mRNA granules under the spine in the early phase after cLTP induction, i.e. at 15’, followed by a translocation of at least the BDNF-Ex1 mRNA variant inside the spine at 30’ and a return to baseline at 60’ from cLTP stimulation.

**Figure 7.**
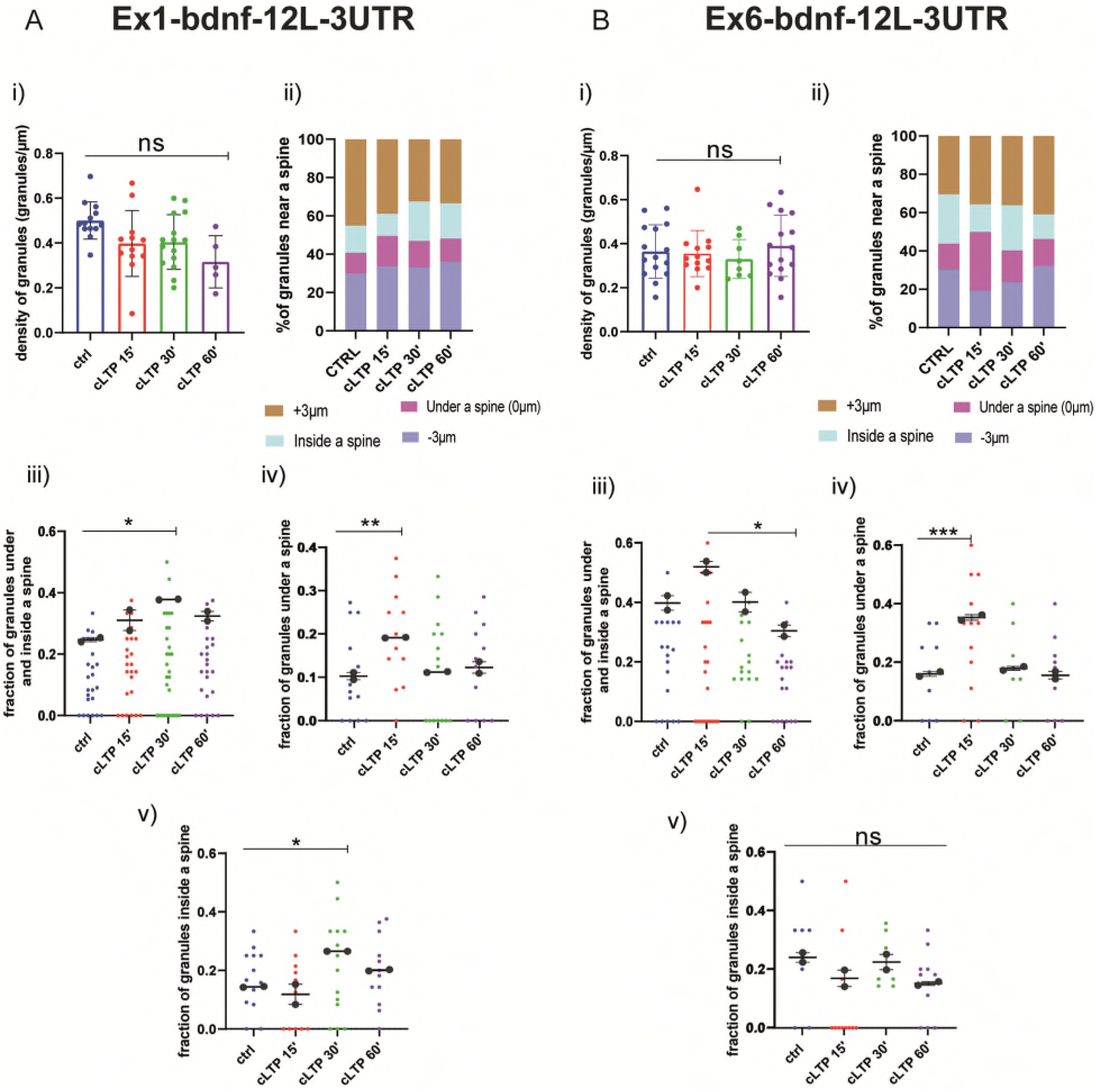
BDNF mRNA granules trafficking near a spine A) Ex1-bdnf-12L-3UTR. i) mRNA granules counted in each dendrite stretches of 30-50 μm ANOVA followed by Dunnett’s post hoc test revealed no significant difference in the density of Ex1-bdnf mRNA granules after cLTP stimulation ii) mRNA granules distribution near a spine after cLTP induction. Granules under a spine and inside a spine were considered iii) Ex1bdnf granules under and inside a spine were considered at different cLTP induction. After 30’ from cLTP induction there is an increase in the granules present under and inside a spine. iv) Ex1bdnf granules under a spine were significantly higher after 15’ from cLTP induction ctrl vs cLTP 15’ p=0.047 v) Ex1bdnf granules inside a spine were significantly higher after 30’ from cLTP ctrl vs 30’ p=0.0253 **B) Ex6-bdnf-12l-3UTR** i) mRNA granules counted in each dendrite stretches of 30-50 μm ANOVA followed by Dunnett’s post hoc test revealed no significant difference in the density of mRNA CamkIIα granules after cLTP stimulation ii) mRNA granules distribution near a spine after cLTP induction. Granules under a spine and inside a spine were considered iii) Ex6bdnf granules under and inside a spine were considered at different cLTP induction. There is a tendency (ctrl vs cLTP 15’ p=0.0785), while there is a significant increase between 15’ and 60’ from cLTP induction iv) Ex6bdnf granules under a spine were significantly higher after 15’ from cLTP induction p= ctrl vs 15’ cLTP p=0.0005 v) Ex6bdnf granules inside a spine were not significantly higher after any time from cLTP. Statistical analysis was done by ANOVA followed by Tukey’s post hoc test.

### 3.6. Basal trafficking and distribution of BDNF protein in living neurons

After having established that cLTP induces a trafficking arrest of BDNF mRNA granules and a redistribution in which they accumulate under the spines and in part inside the spines, it remained to be determined if and how is the BDNF protein redistributed during LTP.

In this series of experiments, the dynamic movements of the BDNF protein were recorded in living neurons thanks to one chimeric construct consisting in the BDNF coding sequence (CDS) common to all BDNF transcript variants fused to the green-fluorescent protein (GFP). It was previously demonstrated that the mRNA derived from this construct, named CDS-BDNF-GFP, has a constitutive dendritic localization and generates a protein which is secreted from the post-synaptic terminals (Baj et al., 2011; Brigadski and Leßmann, 2020; Chiaruttini et al., 2009). Neurons were co-transfected at DIV 9-10 using the CDS-BDNF-GFP construct together with mCherry as a filler and were imaged 2 days later, as previously reported (Baj et al., 2011). As shown in **Figure 8A and B**, neurons transfected with CDS-BDNF-GFP showed clearly identifiable fluorescent protein spots in dendrites (**Figure 8A**) and within one spine (**Figure 8B**). The histogram shown in **Figure 8C** presents the distribution of CDS-BDNF-GFP spots in the dendritic compartment at increasing distance from the soma of neurons cultured in basal conditions. Specifically, at a distance of less than <20 μm from the cell soma only 15 spots were found, 60 spots were counted at 20-60μm from the cell soma, 25 spots at 60-100 μm from the cell soma and finally, 15 spots were detected at a distance further than 100 μm from the cell soma (n=21 neurons; n=3 independent cultures). In untreated cultures, CDS-BDNF-GFP protein spots were detected in both apical (59%) and basal dendrites (41%) (**Figure 8D, left**) and the confined population represented almost the totality of all GFP-tagged BDNF protein spots (99.40%; **Figure 8D, right).**

**Figure 8.**
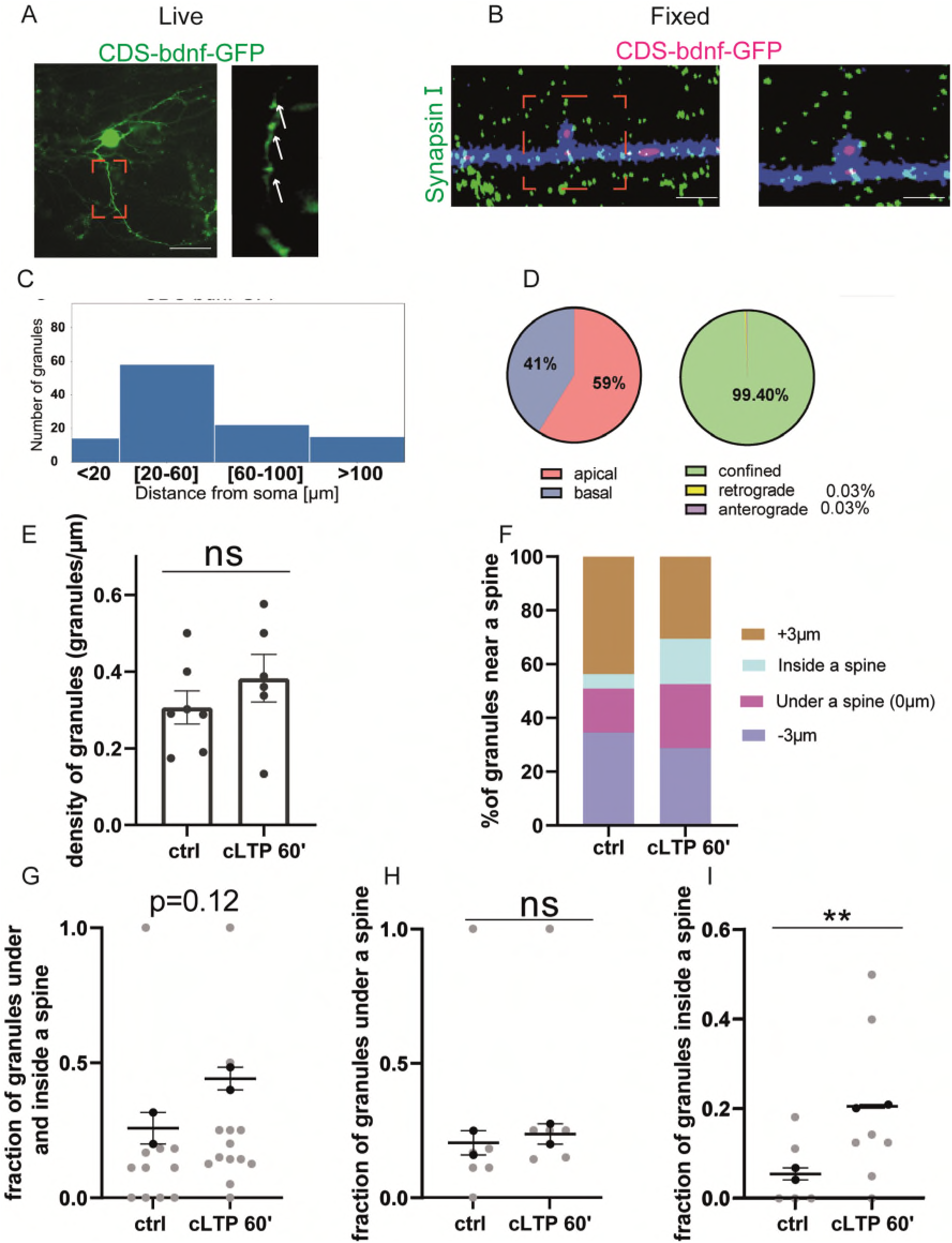
BDNF protein near a spine: A,B) Representative images of CDS-bdnf-GFP spots transfected neurons. A) Living neurons transfected with CDS-bdnf-GFP. B) Neurons were fixed and labelled with Synapsin I. Scale bar 2 μm. C) Histogram of the CDS-bdnf-GFP protein spots distribution in the different dendritic compartments. D) Percentage of CDS-bdnf-GFP spots granules at apical and basal dendrites (left) and the percentage of spots confined, anterograde and retrograde (right). Almost all the protein spots resulted to be confined. E) CDS-bdnf-GFP spots on the left, protein spots counted in each dendrite streches of 30-50 μm. Unpaired T-test revealed no significant difference in the density of CDS-BDNF spots after cLTP stimulation F) protein spots distribution near a spine after cLTP induction. Spots under a spine and inside a spine were considered G) protein spots under and inside a spine were considered at different cLTP induction. After 60’ from cLTP induction there is an tendency p=0.12 H) CDS-BDNF granules under a spine were not significantly higher after cLTP induction I) CDS-BDNF spots inside a spine were significantly higher after 60’ from cLTP ctrl vs 60’ p=0.0085

### 3.7. cLTP-induced redistribution of the BDNF protein in proximity to a spine

The same analysis performed to study the effects of cLTP on mRNA granules distribution and motility, was carried out also for the BDNF protein. Using the fluorescent CDS-BDNF-GFP construct, we found that the density of BDNF protein was not changed after 60 minutes from the induction of cLTP (**Figure 8E**), ctrl vs. cLTP 60’ p= 0.12; n= 8-10 neurons). Of note, during the live imaging we detected newborn spots with a frequency of 8.64% of the total spot population. However, these events seemed to not significantly affect the density of protein spots after 60 minutes from cLTP induction, possibly because of physiological secretion or turnover of the protein. The distribution of the BDNF protein spots near, under or within a spine was analysed (**Figure 8F**, n=15-20 neurons) and a significant increment of BDNF protein spots inside a spine was found at 60’ after cLTP induction (ctrl vs cLTP 60’ p= 0.0085). Taken together, these results suggest that in the late phase after cLTP induction (1h), BDNF becomes more present inside dendritic spines.

## 4. Discussion

The main biological question behind this study concerns the dynamics of BDNF mRNA trafficking and its local translation at synapses in response to synaptic plasticity. Specifically, our study shows that the early phase following LTP induction (i.e. 15’) is characterized by a general arrest of dendritic mRNA trafficking, accompanied by an increase in the local density of mRNA granules below, but not inside the spines. Indeed, the behaviour of the control CaMKIIα mRNA and exon-1 and exon-6 BDNF mRNA was similar in the early-LTP phase, with an increase in the number of mRNA granules under the spine at 15’ and 30’, but it differed in the late phase of LTP (i.e. 60’), in which CaMKIIα mRNA, but not BDNF mRNA, resulted increased also inside the spine. At the protein level, we found that bonafide newly synthesized BDNF was accumulated inside the spine at 60’ after cLTP induction. Taken together, these results suggest that LTP triggers a process involving protein translation at the basis of the spine from locally confined mRNAs, followed by translocation of the protein (or more likely, the vesicles containing it) inside the spines, towards the sites of release at synapses.

A first goal of this study was to study the dynamics of BDNF mRNA and protein trafficking in living neurons. The tool chosen for the visualization of BDNF mRNA was the MS2 stem loops, consisting of 12 bacteriophage loops cloned to our cDNA of interest (ex1-cdsBDNF-3 UTR and ex6-cdsBDNF-3UTR). This is a well established technique that has been widely used before (Tutucci et al., 2017, Holt et al., 2029). Although the actual number of MS2 loops varies from 6 to 120 in the various previous studies (Tutucci et al., 2017; Bauer et al., 2019), we found that 12Loops was sufficient to provide a strong signal that could be reliably recorded in live imaging analysis within a reasonable dynamic range after compensating for the natural bleaching of the signal. For the protein visualization, chimeric BDNF-GFP constructs have been used. Also this tool is very well established for BDNF analysis (Baj et al., 2011; Brigadski and Leßmann, 2020) likely because GFP protein dimerization, a major problem in the use of GFP as a reporter protein, is not relevant in a molecule such as BDNF, which is a dimer by itself. Of note, in order to obtain a more natural behaviour of the mRNA, the two BDNF exon1 and exon6 isoforms used here, included the 3’UTR Long tail which is considered more abundant in dendritically targeted BDNF mRNAs (Vicario et al., 2015).

The performed analysis suggested that the great majority of granules were confined for all the four constructs (EX1-cds BDNF-12L-3’UTR, EX6-cds BDNF-12L-3’UTR, CamkIIα-8L-3’UTR and 12L). This result is in agreement with a recent paper (Donlin-asp et al., 2021) that used molecular beacons to track endogenous CamkIIα and PSD95 mRNA. In analogy with that study, we also found that the vast majority of granules having an anterograde or retrograde movement, stopped and remain confined after the cLTP induction. The soundness of our approach using an exogenous system, as the MS2 stem loop technique, was thus confirmed by the comparison with similar findings (Donlin-Asp et al., 2021) where the endogenous mRNAs were detected by a more physiological method using molecular beacons.

Intriguingly, the EX1-cds BDNF-12L-3’UTR, EX6-cds BDNF12L-3’UTR, CamkIIα-8L-3’UTR showed a slight bias, even though not significant, for anterograde movements. This bias could be caused by the 3’UTR sequence, that, as already reported, is important for the dendritic localization of BDNF mRNA (Vicario et al., 2015) and other mRNAs (Bauer et al., 2019). This explanation is strengthened by the observation that the 12L construct, without the 3’ UTR sequence, did not show any bias for the anterograde movements, showing an equal distribution of anterograde and retrograde movements. The anterograde velocity for the EX6-cds-BDNF-12L-3’UTR showed a significant increment when compared with the anterograde velocity for the EX1-cds BDNF-12L-3’UTR. This can be linked to a previously reported finding that the Ex6-BDNF is located at more distal part of dendrites in comparison with the Ex1-BDNF (Baj et al., 2011).

After LTP induction, the large majority of granules became confined. The increment of confined movements was evident for all four constructs, even for the 12L. Although it has been previously reported that the MS2 loops are able to generate RNA granules that can travel along the dendrites (Bauer et al., 2019), it was reasonable to expect that unlike BDNF and CamkII, the 12L construct should not give rise to any biological response to cLTP. Considering 20 neurons imaged for each construct, the total number of granules detected for the 12L was significantly lower than that detected for the other constructs. In addition, the RNA granules formed by the 12L construct showed more proximal localization in comparison with all other constructs used. Furthermore, after cLTP induction the number of anterograde movements was decreased for the EX6-cds BDNF-12L-3’UTR and EX1-cds BDNF-12L-3’UTR, but not for 12L. On the contrary, the number of retrograde movements decreased after cLTP for 12L but not for the BDNF mRNA. Thus, while the movements of BDNF mRNAs appeared to be dictated by cLTP induction, the 12L resulted to be not responsive to it, supporting the lack of any biological significance for this construct in neurons.

Previous studies showed that Ex1 BDNF transcript was mainly located at proximal dendrites, whereas Ex6 BDNF mRNA has been identified beyond 100 µm from the cell soma (Chiaruttini et al., 2009; Baj et al., 2011; Mallei et al., 2015). Surprisingly, upon stimulation with Forskolin, the EX6-bdnfCDS-12L3UTR could not be detected in the most distal dendritic compartment. This could be most likely due to the timing and nature of the stimulus. Indeed, using a similar constructs with the 12L replaced by the GFP (EX6-bdnfCDS-GFP-3UTR) a very distal localization was achieved after 3 hours of stimulation with 10 mM KCl (Baj et al., 2011) while in this study, 15 minutes of Forskolin 10 µM might have not been enough for BDNF mRNAs to reach the distal compartment of the dendrites.

One further goal of this study was to understand if after cLTP induction, BDNF mRNA dynamically accumulates and whether it is locally translated into BDNF protein in proximity of spines or if mRNA and protein have a random distribution along the dendrites. Using super-resolution microscopy, here we measured the distance between a granule and the nearest spine and whether the granules were either below or inside the spine. Although with this analysis it was not possible to specifically identify the potentiated spines, results highlighted that in the early-LTP phase, both CamkIIα and BDNF granules were generally increased under the spines, while in the late-LTP phase only CamkIIα granules were accumulated inside the spine. A possible interpretation is that after short periods (15-30 minutes) from cLTP induction, a rise in Ca^2+^ concentrations in the dendritic shaft underneath activated spines provokes an uncoupling of the mRNA granules from the kinesins/microtubule-driven transport leading the mRNAs to become stationary near the activated spines. While for BDNF mRNA granules the dendritic segment under the spine is the final destination, likely due to the need of this secreted protein to be processed through the secretory pathway, CamkIIα mRNA which encodes for a cytoplasmatic protein become further recruited by an actin-based transport system which drives this mRNA into the spines where it becomes locally translated (Hanus and Ehlers, 2008). This model is compatible, with the finding that there was an increment of BDNF protein spots inside a spine after 60 minutes from cLTP induction. Of note, the construct with only the 12 stem loops (12L) did not show any significant increase in the number of granules under or inside the spine, confirming that the distribution of the 12L granules is cLTP independent.

Taking into consideration that local BDNF synthesis plays a critical role in L-LTP, then the dendritic localization of all the machinery for the translation of secretory protein is necessary. There are indeed a number of post-translational modifications for BDNF, including glycosylation, proper folding, cleavage and sorting to constitutive or regulated secretion pathway. Folding and N-glycosylation are processed in ER whereas cleavage and sorting occur in Golgi apparatus. In the past years, Ehlers and colleagues demonstrated that at least in cultured hippocampal neurons, Golgi apparatus are absent in majority of dendrites (Wang et al., 2012; Horton and Ehlers, 2003). However, it has been showed that small Golgi-like organelles (so-called Golgi outposts) are selectively localized to dendritic branch points. Thus all the secretory pathway is available in dendrites. However, in a previous study using electron microscopy in situ hybridization, we found that BDNF mRNA could be detected in association with polyribosomes located under spines and near dendritic branchings, suggesting active translation without interaction with the endoplasmic reticulum (Tongiorgi et al., 2004). At present, how locally synthesized secretory or transmembrane proteins which are translated on polyribosome can get processed through the secretory route remains a conceptually challenging issue.

## 5. Acknowledgements

This study was supported by a grant from the Italian Ministry of University Education and Research (MIUR) PRIN 2017 Prot. 2017HPTFFC “SYNACTIVE: Synaptic engrams in memory formation and recall”. GB was supported by a doctoral fellowship from Fondo Sociale Europeo (FSE) of the Regione Friuli Venezia Giulia. The authors are thankful to Gabriele Baj and Agnès Thalhammer from the Centro Interdipartimentale di Microscopia Avanzata (CIMA) of the University of Trieste, for valuable discussions and technical advice on microscopy.

